# Restoration of motor-evoked cortical activity is a distinguishing feature of the most effective rehabilitation therapy after stroke

**DOI:** 10.1101/2020.03.05.974972

**Authors:** Emilia Conti, Anna Letizia Allegra Mascaro, Alessandro Scaglione, Giuseppe de Vito, Francesco Calugi, Maria Pasquini, Tommaso Pizzorusso, Silvestro Micera, Francesco Saverio Pavone

## Abstract

**Background:** An ischemic stroke is followed by the remapping of motor representation and extensive changes in cortical excitability involving both hemispheres. Although stimulation of the ipsilesional motor cortex, especially when paired with motor training, facilitates plasticity and functional restoration, the mechanisms underneath the reshaping of cortical functionality are widely unknown.

**Objective:** We investigated the spatio-temporal features of motor-evoked cortical activity associated with generalized recovery after stroke, and its dependence on the type of rehabilitative treatment.

**Methods:** We designed a novel rehabilitative treatment that combines neuro-plasticizing intervention with motor training. Specifically, optogenetic stimulation of peri-infarct excitatory neurons expressing Channelrhodopsin-2 was associated with daily motor training on a robotic device. The effectiveness of the combined therapy was compared with spontaneous recovery and with the single treatments (i.e. individually administered optogenetic stimulation or motor training).

**Results:** We found that only the combined therapy promotes generalized recovery of forelimb function and the rescue of spatio-temporal features of motor-evoked activity. Generalized recovery results from a new excitatory/inhibitory balance between hemispheres as revealed by the augmented motor response flanked by the increased expression of parvalbumin positive neurons in the peri-infarct area.

**Conclusions:** Our findings demonstrate that though behavioral recovery is not necessarily associated with the restoration of pre-stroke motor-evoked activity, the reestablishment of pre-stroke activation transients was a distinguishing feature of the most efficient therapeutic approach, the combined therapy.

## Introduction

Ischemic injuries within the motor cortex result in functional deficits that may profoundly alter patients’ quality of life. Survivors are often chronically impaired with long-term disability^1^. To regain sensory and motor functions after stroke, spared neural circuits must reorganize^2–4^. Multiple strategies were developed to enhance neural rewiring which dramatically improved functional recovery^4^ including pharmacological treatment, motor training, and brain stimulation. Among them, cortical neuromodulation techniques, such as transcranial magnetic stimulation (TMS) and transcranial direct current stimulation (tDCS), represent a promising non-invasive approach to improve cortical remapping. Nevertheless, these treatments can induce diffuse and non-specific activation in mixed neuronal populations^5–7^, revealing the necessity for more targeted therapies. With the emergence of optogenetics, specific neuronal populations can be activated or inhibited achieving high temporal and spatial precision^8–11^. Recently, optogenetics has been proficiently used to selectively modulate the excitatory/inhibitory balance of brain circuits affected by a stroke lesion^12,13^. Repeated optogenetic neuronal stimulation of the ipsilesional hemisphere induced a significant improvement in neurovascular coupling response^14^. Furthermore, chronic optogenetic stimulation of the entire cortical mantle promoted behavioral recovery associated with the formation of new and stable thalamocortical synaptic boutons^15^. Nevertheless, information on the remapping of motor representation and motor-evoked cortical activation following optogenetic stimulation is largely unexplored. Further, no study investigated how stimulation-induced cortical remapping correlates with functional recovery.

When combined with motor training, cortical stimulation creates a pro-plasticizing milieu where spared neurons are more susceptible to experience-dependent modifications^4,16,17^. Though several studies investigated the effect of combining neuronal modulation, such as TMS^18–20^ and tDCS^21–23^, with robotic training, results are still contradictory. Up to now, whether the combination of ipsilesional neuronal stimulation and physical training promotes motor recovery is still unknown. Further, though it is established that the combination of neuronal modulation and motor training plays a key role in post-stroke recovery, no investigation has yet addressed if possible physical improvement is supported by alterations in motor maps and in the distributed motor-evoked cortical activation.

Our group and others recently demonstrated that optical tools, including *in vivo* fluorescence imaging and optogenetics, are a useful mean to characterize critical features of plasticity, repair and recovery after stroke^24–29^.

Here, we designed a light-based stimulation protocol of peri-infarct excitatory neurons as a rehabilitative approach to achieve generalized recovery. To dissect cortical remapping in motor-evoked neuronal activity in peri-infarct area we took advantage of wide-field fluorescence imaging over the affected hemisphere. We found that longitudinal optogenetic stimulation restored forelimb functionality but motor-evoked functional activity was not recovered. However, coupling optogenetic stimulation with longitudinal motor training of the impaired forelimb on a robotic platform halved the time required for a full recovery of forelimb function compared to optogenetic stimulation only. Furthermore, the rapid behavioral recovery was associated with the restoration of temporal features of calcium transient such as peak amplitude and slope. The analysis of motor-evoked activation maps in double treated mice identified the peri-infarct area as the region of the cortex mostly involved in motor task. Finally, the combined treatment promoted the restoration of an interhemispheric balance between the two hemispheres, revealed by an increase of expression of Parvalbumin positive cells in the peri-infarct area, and plasticity marker, GAP43, both in per-infarct neurons and in contralesional hemisphere’s fibers.

## Methods

### Mice

All procedures involving mice were performed in accordance with regulations of the Italian Ministry of Health authorization n. 871/2018. Mice were housed in clear plastic cages under a 12h light/dark cycle and were given ad libitum access to water and food. We used a transgenic mouse line, C57BL/6J-Tg(Thy1GCaMP6f)GP5.17Dkim/J, from Jackson Laboratories (Bar Harbor, Maine USA). Mice were identified by earmarks and numbered accordingly. Animals were randomly divided into 5 groups. Each group contained comparable numbers of male and female mice (weighing approximately 25g). Age of mice (ranging from 6 to 8 months old) was consistent between groups.

### Experimental design

Animals were distributed in 5 groups as follows: Sham n=6; Stroke n=4; Optostim n=4; Robot= 7; Optostim+Robot (abbreviated in OR) n= 4 (Supplementary Figure 2).

- The Sham group consisted of 6 healthy mice. During surgery instead of induced photothrombosis we intraperitoneally injected saline and then we illuminated the primary motor cortex (+1.75 ML and +0.50 AP). We then intracranially injected saline in the sensory cortex (+1.75 ML and −0.75 AP). After 5 days of recovery from surgery mice performed 5 days of motor assessment on the robotic platform in order to investigate cortical activation during a motor task in healthy conditions.
- The Stroke group consisted of 4 mice. At the beginning of the protocol we evaluated forelimbs use via Schallert test. We induced a focal stroke in the primary motor cortex (+1.75 ML and +0.50 AP). During the same surgery we intracranially injected saline in the sensory cortex (+1.75 ML and −0.75 AP). Two days after surgery we performed behavioral tests to identify alteration in forelimb use consequent to stroke. We performed behavioral experiments at the end of each week to longitudinally investigate mice spontaneous recovery. After 5 days of recovery from surgery we stimulate Stroke mice, though not expressing Channelrhodopsin-2 (ChR2), with a blue laser for 20 days to evaluate possible artifacts due to repeated laser stimulation. After 25 days of spontaneous recovery mice performed 5 days of motor assessment on the robotic platform in order to investigate motor-evoked cortical activation. 30 days after stroke mice were perfused.
- The Optostim group consisted of 4 mice. At the beginning of the protocol we evaluated forelimbs use via Schallert test. We induced a focal stroke in the primary motor cortex. During the same surgery we intracranially injected an adeno associated virus (AAV9-CaMKII-ChR2-mCherry) to induce the expression of ChR2 in the sensory areas. Two days after surgery we performed behavioral tests to identify alteration in forelimb use consequent to stroke. After 5 days of recovery from surgery mice began the rehabilitation paradigm consisting in 20 days of optogenetic stimulation of the peri-infarct area. 25 days after photothrombosis, mice performed 5 days of motor assessment on the robotic platform in order to investigate motor-evoked cortical activation. At the end of each week of rehabilitation we performed behavioral experiments to longitudinally investigate mice recovery. 30 days after stroke mice were perfused.
- The Robot group consisted of 7 mice. At the beginning of the protocol we evaluated forelimbs use via Schallert test. We induced a focal stroke in the primary motor cortex. During the same surgery we intracranially injected an adeno associated virus (AAV9-CaMKII-ChR2-mCherry) to induce the expression of ChR2 in the sensory areas. Two days after surgery we performed behavioral tests to identify alteration in forelimb use consequent to stroke. After 5 days of recovery from surgery mice began the rehabilitation paradigm consisting in 20 days of robotic training. At the end of each week of rehabilitation we performed behavioral experiments to longitudinally investigate mice recovery. 30 days after stroke mice were perfused.
- The Optostim+Robot (OR) group consisted of 4 mice. At the beginning of the protocol we evaluated forelimbs use via Schallert test. We induced a focal stroke in the primary motor cortex. During the same surgery we intracranially injected an adeno associated virus (AAV9-CaMKII-ChR2-mCherry) to induce the expression of ChR2 in the sensory areas. Two days after surgery we performed behavioral tests to identify alteration in forelimb use consequent to stroke. At the end of each week of rehabilitation we performed behavioral experiments to longitudinally investigate mice recovery. After 5 days of recovery from surgery mice began the rehabilitation paradigm consisting in 20 days of robotic training followed by optogenetic stimulation of the peri-infarct area. 30 days after stroke mice were perfused.

### Surgical procedures

Mice were injected with a Rose Bengal solution (0.2 ml, 10 mg/ml solution in Phosphate Buffer Saline (PBS)). Five minutes after intraperitoneal injection a white light from an LED lamp was focused with a 20X objective to illuminate the primary motor cortex (M1) for 15 min inducing unilateral stroke in the right hemisphere. During the same procedure, we delivered 0.5 μl of AAV9-CaMKIIa-hChR2(H134R)-mCherry (2.48*10^13^ GC/mL) 600μm deep inside the cortex at −0.75 anteroposterior, +1.75 mediolateral. For further information see Supplemental Material.

### Robotic Platform

Animals were trained by means of the M-Platform^30,31^, which is a robotic system that allows mice to perform a retraction movement of their left forelimb. Motor rehabilitation consists in a pulling task: first animal forelimb is passively extended by the linear actuator of the platform and then the animal has to pull back forelimb up to the resting position. Motor training is composed by 15 movements and after each movement the animal receives a liquid reward. For further information see Supplemental Material.

### Optogenetic stimulation

Daily optogenetic stimulation was performed on head-fixed awake mice by employing a 473 nm laser delivering 5 Hz, 10ms light pulses. Laser power, ranging from 0.2 to 0.8 mW, was daily adjusted according to the increment of the transfected area and the progressive lowering of stimulation threshold over the weeks. For further information see Supplemental Material.

### Statistical analysis

Statistical analysis was performed using OriginPro software (OriginLab Corporation) ans results were considered statistically significant with a p value <=0.05. For the Schallert test analysis, if different groups were compared at the same time point, a two-way repeated measure ANOVA with factors GROUP and TIME was used. In case where different timepoints of the same group were compared, a one-way ANOVA with factor GROUP was used.

For calcium, forces and immunohistochemical analysis, a one-way ANOVA was used, with factor GROUP. For all ANOVAs that were statistically significant, multiple comparison among groups, or time points, were assessed using Tukey HSD test.

For information regarding Schallert test, wide-field microscope, and image, forces, and immunohistochemical analysis see Supplemental Material.

## Results

### Peri-infarct optogenetic stimulation restores forelimb function but not cortical activation features

The main goal of this study is to find a neuronal substrate of generalized functional recovery within the distributed motor-evoked cortical activity. To this aim, we compared behavioral and calcium imaging data from mice receiving three different treatments, optogenetic stimulation, motor training and a combination of them. First, the efficacy of repeated optogenetic stimulation of peri-infarct excitatory neurons was tested in stroke mice. A photothrombotic stroke was induced on the M1 of the right hemisphere in Thy1-GCaMP6f mice (Supplementary Figure 1A). We developed a rehabilitation protocol based on longitudinal optogenetic stimulation of peri-infarct excitatory neurons expressing Channelrhodopsin-2 (ChR2, Figure 1A and Supplementary Figure 1B-D). Optogenetic stimulation was performed daily and consisted of three successive 30-sec laser stimulation trains, separated by 1-min rest intervals (Figure 1B). The optogenetic therapy lasted 4 weeks starting 5 days after stroke. We divided our sample into 3 groups (Figure 2A, Supplementary Figure 2 and Experimental design Materials and Methods section): Sham (healthy mice, no stroke), Stroke (stroke, spontaneous recovery), Optostim (stroke, optogenetic rehabilitation). During the last week of our investigation mice performed 5 days of motor assessment in order to evaluate motor-evoked cortical activity (Figure 1C).

**Figure 1.**
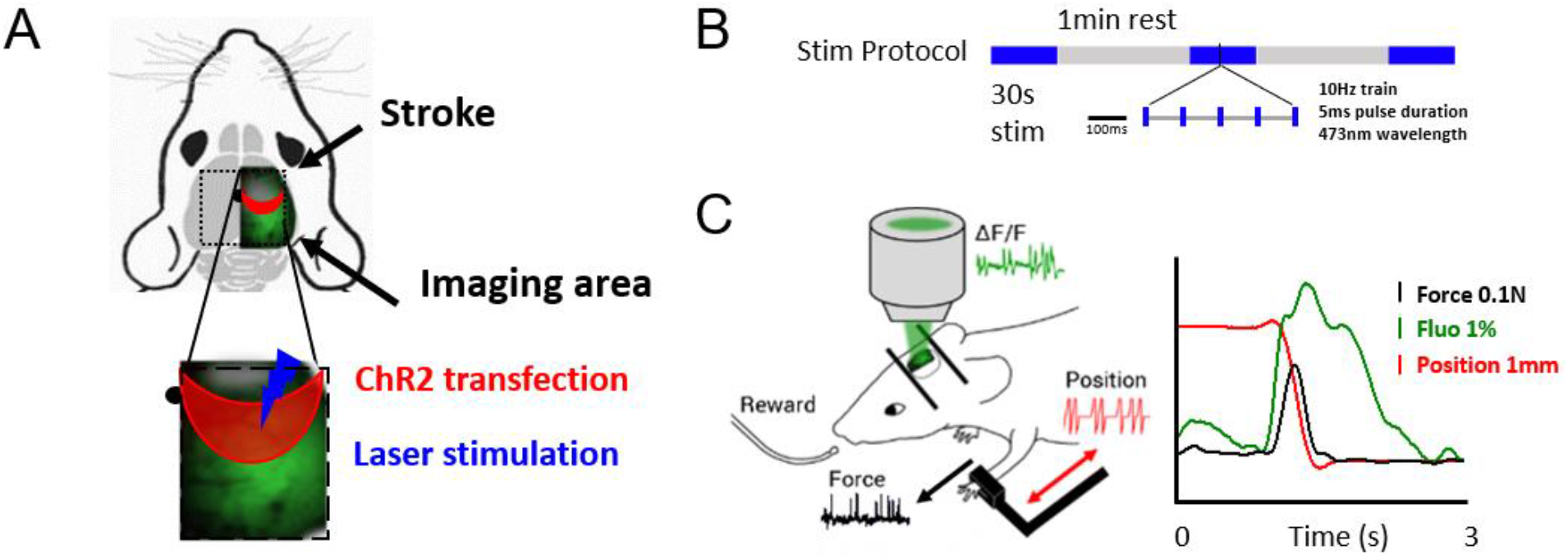
Experimental design: (A) Field of view graphical representation: ChR2-expressing neurons (red area) are stimulated by blue laser. Cortical activity is revealed in the right hemisphere during motor assessment. Grey cloud represents stroke core, black dot represents bregma. (B) Stimulation paradigm consisting of 3 stimulation train separated by 1-min rest intervals. (C) Graphical representation of M-Platform for motor assessment. Graph on the right shows an overlap of simultaneously recorded traces (Force, Fluorescence and Position) of an exemplary retraction movement.

**Figure 2.**
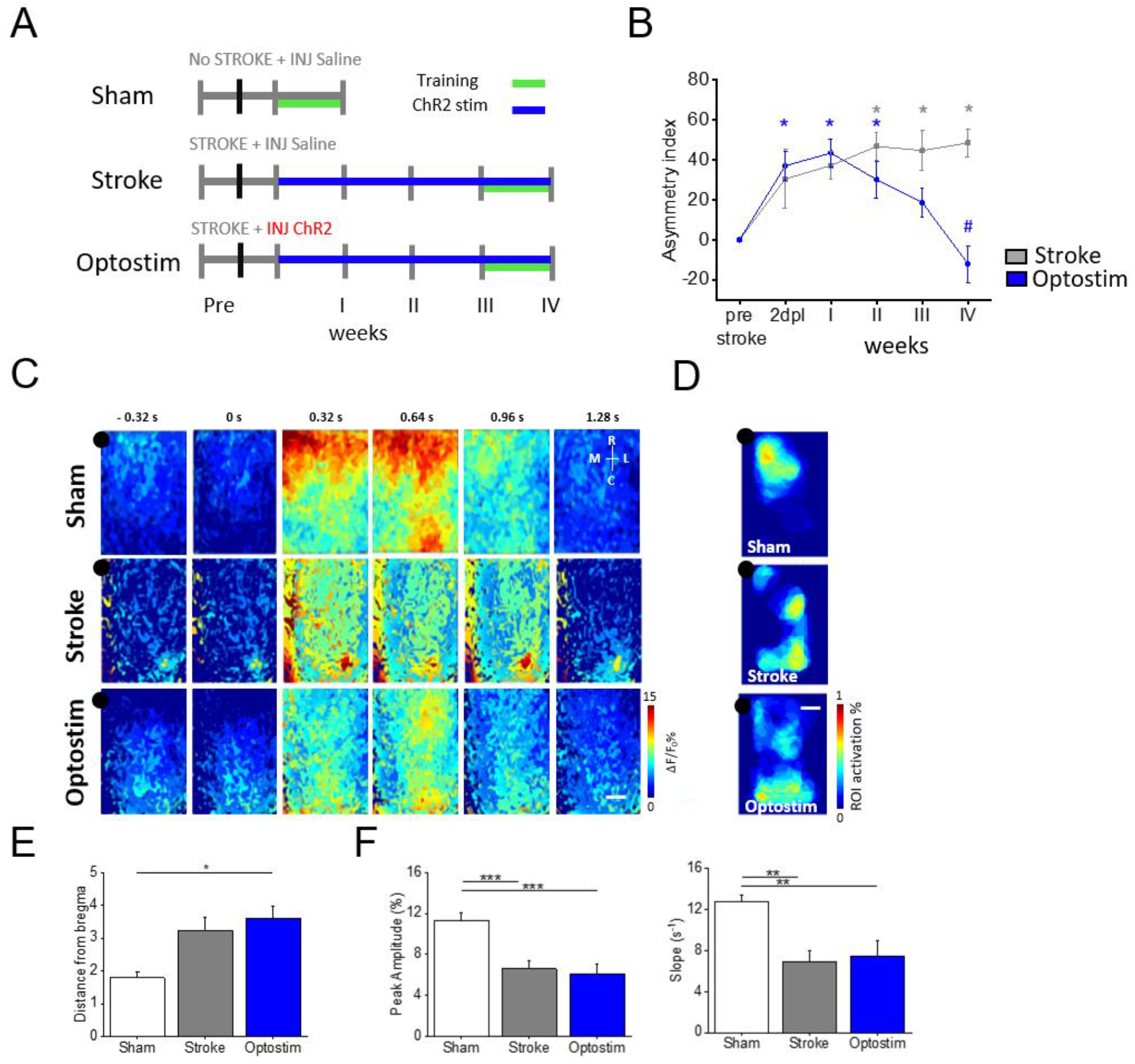
Optogenetic stimulation of peri-infarct area promotes the recovery of forelimb functionality but not the restoration of spatio-temporal cortical profiles: (A) Sham, Stroke and Optostim experimental timeline. (B) Pre- and post-lesion performance of Stroke (Grey) and Optostim (Blue) groups measured as Asymmetry Index in the Schallert Cylinder test. *p<0.05 based on two-way ANOVA repeated measure followed by Tukey’s correction refers to comparison of post-stroke time points in each group designated by color to respective pre-stroke conditions. Stroke group: p IIw = 0.014; p IIIw = 0.020; p IVw = 0.010; Optostim group: p 2dpl = 0.037; p Iw = 0.001; p IIw = 0.038. #p<0.05 refers to comparison between Optostim and Stroke groups. Optostim vs Stroke pIVw <0.0001 (C) Image sequence of cortical activation assessed by calcium imaging during pulling of contralateral forelimb, from 0.32 s before to 1.28 s after the onset of the force peak. Each row shows a representative sequence from a single animal for each group. Scale bar 1mm. (D) Motor-evoked activation maps show the average thresholded region of maximum activation triggered by the pulling task for each experimental group. Scale bar 1mm. (E) Graph shows the distance from bregma (average ± SEM) of maps centroid (Sham = 1.8 ± 0.2 mm; Stroke = 3.2 ± 0.4 mm; Optostim = 3.6 ± 0.4 mm; *p<0.05 based on one-way ANOVA followed by Tukey’s correction: pSham-Optostim =0.03). (F) Left panel: graph shows the maximum of fluorescence peaks (average ± SEM) of calcium transient (Peak-Amplitude_Sham = 11.3 ± 0.7%; Peak-Amplitude_Stroke = 6.6 ± 0.7%; Peak-Amplitude_Optostim = 6.1 ± 0.9%; ***p<0.0005 based on one-way ANOVA followed by Tukey’s correction: pSham-Stroke= 0.0005; pSham-Optostim =0.0002). Right panel: graph shows the slope (average ± SEM) of the rising phase of fluorescence traces (Slope_Sham = 12.8 ± 0.6 s^-1^; Slope_Stroke = 6.9 ± 1.0 s^-1^, Slope_Optostim = 7.4 ± 1.5 s^-1^; **p<0.005 based on one-way ANOVA followed by Tukey’s correction: pSham-Stroke = 0.002; pSham-Optostim = 0.005). nSham=6; nStroke=4; nOptostim=4.

We longitudinally estimated generalized recovery of forelimb functionality by performing Schallert test at the end of each week of treatment. While the asymmetry index (Ai) in spontaneously recovered mice (Stroke group) did not reach pre-stroke levels, a full recovery of forelimb function was achieved after 4 weeks of daily optogenetic stimulation (Optostim group, Figure 2B).

We then assessed if generalized behavioral recovery was associated with specific features of motor-evoked cortical functionality. To this aim, we performed wide-field calcium imaging of the affected hemisphere on GCaMP6f mice during the execution of a pulling task within a robotic device, the M-Platform^30,31^. The M-Platform was previously integrated with a custom-made wide-field mesoscope to perform calcium imaging of motor-evoked activity and optogenetic stimulation^32–34^. This robotic device allowed a detailed assessment of motor-evoked cortical activation during an active forelimb pulling task.

To evaluate possible spurious activation of ChR2 expressing neurons due to the blue LED for calcium imaging, we performed control experiments imaging activity in Thy-GCaMP6f mice with or without ChR2 injection while performing 4 weeks of motor training. No significant difference was visible in calcium transients from ChR2^+^ mice as compared to ChR2^-^ mice (Supplementary Figure 3A), demonstrating that ChR2 stimulation induced by the imaging LED was negligible or absent.

We then analyzed the spatial extension of motor representation in the ipsilesional hemisphere (Supplementary Figure 3B) by overlapping the movement-triggered activation maps obtained for each day of training. This comparison showed a segregated activation in healthy mice, whereas in Stroke and Optostim mice the maps covered most of the affected hemisphere, up to the caudal regions of the cortex (such as retrosplenial and visual areas Figure 2C-D), as quantified by analyzing distance from bregma of motor-evoked map’s centroid (Figure 2E). Then, we explored the possibility that temporal features of activation might be correlated with generalized recovery in the Optostim group. However, by analyzing the fluorescence transients averaged from the region of maximum calcium activation during active pulling, we observed no significant differences in amplitude and timing in Optostim mice compared to spontaneously recovering Stroke mice (Figure 2F). These results demonstrate that, even though recovery of forelimb function can be achieved by optogenetic stimulation alone, the spatiotemporal features of motor-evoked cortical activity in the ipsilesional hemisphere did not recover to pre-stroke conditions.

### Combining optogenetic stimulation with motor training boosts the generalized recovery and is associated with reshaping of motor-evoked activation maps and restoration of temporal features of calcium transients

In a previous study, we showed that a combined rehabilitation protocol of pharmacological inactivation and motor training was beneficial to achieve a full generalized recovery^29^. We wondered if an even more effective recovery could be achieved by coupling optogenetic stimulation to longitudinal motor training of the affected forelimb on the M-platform. Thus, a rehabilitation protocol combining daily training on the M-Platform and optogenetic stimulation was tested (OR group, Figure 3A and Supplementary Figure 2). We choose to sequentially perform motor training and optogenetic stimulation of the peri-infarct cortex to avoid a potential bias on fluorescence signal during motor training (calcium imaging) due to the previous excitation of the cortex. The combined treatment was applied daily from the acute phase (5 days after stroke) up to 4 weeks after stroke. The OR group was then compared to healthy mice (Sham group), spontaneous recovery (Stroke group), and motor exercise alone (Robot group).

**Figure 3.**
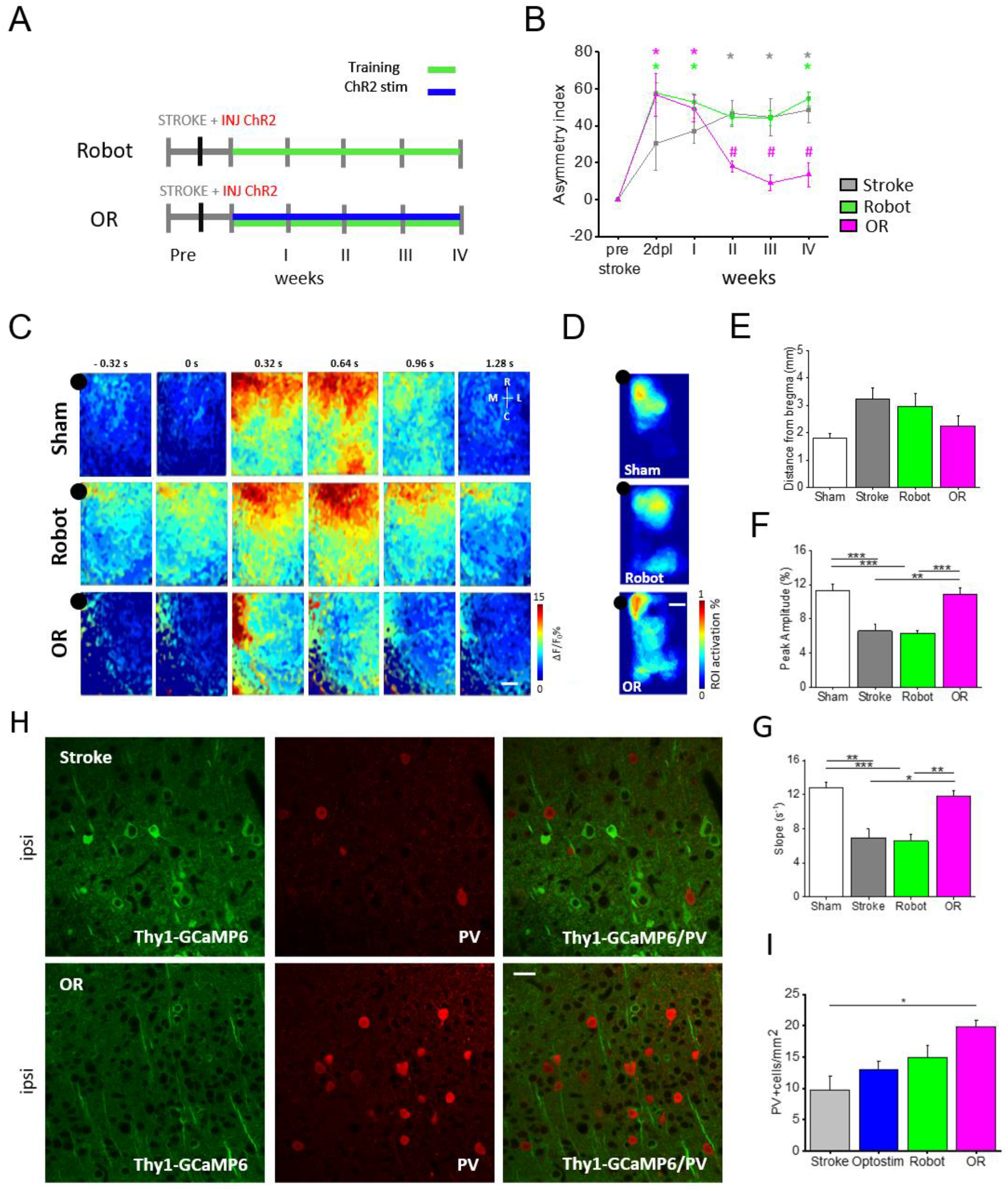
Combining optogenetic stimulation with motor training boosts generalized recovery and promotes the restoration of motor-evoked cortical functionality: (A) Experimental timeline for Robot and OR groups. (B) Pre- and post-lesion performance of Stroke (Grey), Robot (Green), and Optostim+Robot (Magenta) groups measured as Asymmetry Index in the Schallert Cylinder test. *p<0.05 based on two-way ANOVA repeated measure followed by Tukey’s correction comparing post-stroke time points in each group designated by color to respective pre-stroke values. Stroke group: p IIw = 0.014; p IIIw = 0.020; p IVw = 0.010; Robot: p 2dpl =0.013; p Iw = 0.021; p IVw = 0.008; Optostim+Robot: p 2dpl < 0.001; Iw <0.001. #p<0.05 refers to comparison between OR and Stroke groups. OR vs Stroke pwII=0,005; pwIII=0,0004; pwIV=0,0006. nStroke=4; nRobot=7; nOR=4. (C) Image sequence of cortical activation during pulling of the handle, from 0.32s before to 1.28s after force peak onset. Each row shows a representative sequence from a single animal of each group Sham (repeated from Figure 1), Robot and OR. Scale bar 1mm. (D) The panel shows the average thresholded ROI computed for each experimental group. Black dot represents bregma. Scale bar 1mm. (E) Graph shows the distance from bregma (average ± SEM) of maps centroid (Sham = 1.8 ± 0.2 mm; Stroke = 3.2 ± 0.4 mm; Robot = 2.9 ± 0.5 mm; OR = 2.2 ± 0.4 mm). (F) The graph shows the maximum of fluorescence peaks (average ± SEM) of calcium transient (Peak-Amplitude_Sham = 11.3 ± 0.7%; Peak-Amplitude_Stroke = 6.6 ± 0.7%; Peak-Amplitude_Robot = 6.3 ± 0.3%; Peak-Amplitude_OR = 10.9 ± 0.7%; **p<0.005, ***p<0.0005 based on one-way ANOVA followed by Tukey’s correction: pSham-Robot = 0.00005; pOR-Stroke= 0.003; pOR-Robot=0.0005). (G) The graph shows the slope (average ± SEM) of the rising phase of fluorescence traces (Slope_Sham = 12.8 ± 0.6 s^-1^; Slope_Stroke = 6.9 ± 2.0 s^-1^, Slope_Robot = 6.5 ± 0.8 s^-1^; Slope_OR = 11.8 ± 0.7 s^-1^; *p<0.05, **p<0.005 ***p<0.0005 based on one-way ANOVA followed by Tukey’s correction: pSham-Robot= 0.0002; p OR-Stroke = 0.02; p OR-Robot=0.004); (H) Transgenic expression of GCaMP6f under Thy1 promoter (green) and representative PV immunostaining (red) of a Stroke and OR mouse. (I) Quantification of PV+ cells in Stroke and OR groups in the peri-infarct area (Stroke= 19,7 ± 5,0; OR= 37,7 ± 8,3; *p<0.05 based on one-way ANOVA followed by Tukey’s correction: pStroke-OR= 0,04). Scale bar 20μm.; nSham=6; nStroke=4; nRobot=7; nOR=4.

Training induced a progressive modulation of the force transients measured on the M-Platform during active pulling of the affected forelimb. A small reduction in amplitude and full-width half maximum (FWHM) was seen both in OR and Robot mice (Supplementary Figure 3C). In addition, the progressive decrease of time-to-target visible in double-treated mice was in line with previous findings in robot-treated stroke mice^31^. We then evaluated alterations in spontaneous forelimbs use on Schallert test. In accordance with Spalletti and colleagues^35^, motor training alone was not able to restore pre-stroke performances (Figure 3B). Indeed by comparing Ai of Stroke and Robot mice no significant differences were revealed between groups. Conversely, OR mice recovered generalized forelimb functionality already on the second week of rehabilitation (Figure 3B), thus supporting the hypothesis that combined rehabilitation boosts recovery. Indeed, as emerges from the comparison with pre-stroke condition the forelimb use is shifted towards the non-paretic limb in the acute phase after stroke (2days post lesion and after I week of treatment). During the rehabilitation period (II-IV weeks of treatment) the asymmetry index was recovered in OR mice to pre-stroke levels, significantly different from spontaneously recovered mice.

We wondered if this fast behavioral improvement due to the combined treatment was mirrored into specific spatiotemporal features of motor-evoked cortical activity. Examples of temporal sequences of pulling-evoked cortical activation are shown in Figure 3C, together with the associated motor representations (i.e. the thresholded motor-evoked maps, see Supplementary Materials and Methods section) on the fourth weeks after stroke (Figure 3D). The activation maps of both Robot and OR groups show a partial recovery to healthy controls (Sham) in terms of distance to bregma, meaning that daily motor training promotes the refocusing of motor representation in the peri-infarct area (Figure 3E). Interestingly, combined training shifts the motor representation in the rostro-medial direction, suggesting that during the last week of rehabilitation the caudal part of the secondary motor cortex was the principal region activated during contralateral forelimb pulling. We further investigated how the temporal profiles of calcium transients averaged over the motor representation area were altered after combined rehabilitation. Amplitude and slope in mice treated with motor training alone (Robot group) were comparable to spontaneous recovery (Stroke group). Conversely, the synergic effect of combined rehabilitation promoted the reestablishment of these salient features of calcium transient in peri-infarct area (Figure 3F, G). This positive trend towards pre-stroke conditions (Sham group) started from the second week of rehabilitation in OR mice (Supplementary Figure 3D). The pronounced increment in peak amplitude and slope of calcium transient in combined rehabilitated mice became comparable to healthy condition after 4 weeks of combined treatment.

The modulation of cortical motor-evoked activity in the various groups could be associated with a new excitatory/inhibitory balance^14^ potentially compensating detrimental consequences of stroke^36^. We thus evaluated the density of parvalbumin-positive (PV^+^) cells throughout all cortical layers of the peri- and contra-lesional cortices (Figure 3H). In the peri-infarct region, a small but not significant increase of PV^+^ cell density was induced by the single treatments (Optostim and Robot, Figure 3I) with respect to spontaneous recovery mice (Stroke group). Interestingly, the PV^+^ cell density is significantly higher in peri-infarct cortex of double treated mice (OR group) compared to non-treated (Stroke) animals (Figure 3I). Conversely, no significant differences were observed in the contralesional hemisphere in all groups (Supplementary Figure 4A). Together with the increased motor-evoked activation levels in pyramidal cells, these results suggest that the synergic effect of combined rehabilitation could promote the establishment of a new excitatory/inhibitory equilibrium in the peri-infarct and contralesional hemispheres.

### Generalized recovery is associated with an increased expression of GAP43

To further identify molecular targets associated with generalized recovery, we tested the presence of plasticity markers in the lesioned and contralesional cortex by immunohistochemical analysis. We examined the expression of the growth-associated protein 43 (GAP43), a plasticity marker involved in synaptic turnover and reorganization after stroke^3,35,37^. While after optogenetic stimulation (Optostim group) or motor training (Robot group) GAP43 expression levels were comparable to spontaneously recovered mice (Stroke group), combined rehabilitative treatment promoted a massive expression of this neuronal plasticity marker in the peri-infarct region (Figure 4A). Though in the contralesional hemisphere no differences in GAP43^+^ cells were observed between groups (Supplementary Figure 4B), an increase in number and length of GAP43^+^ fibers was present in the homotopic areas on the contralesional hemisphere after all treatments (Optostim, Robot, and OR groups; Figure 4B). Nevertheless, only the combined treatment (OR group) showed significantly different density of GAP43^+^ neurites compared to spontaneously recovered mice (Stroke group). This result demonstrate that perilesional stimulation and motor training synergistically enhance the density of GAP43^+^ neurites in distal regions functionally related to the stroke core, which is possibly associated to axonal sprouting^38–40^ or dendritogenesis^41,42^. Taken together, the immunohistochemical analysis shows that generalized behavioral recovery and the associated cortical remapping induced by combined rehabilitation are supported by a plasticizing milieu promoted by the synergic effect of ipsilesional neuronal stimulation and repetitive motor training.

**Figure 4.**
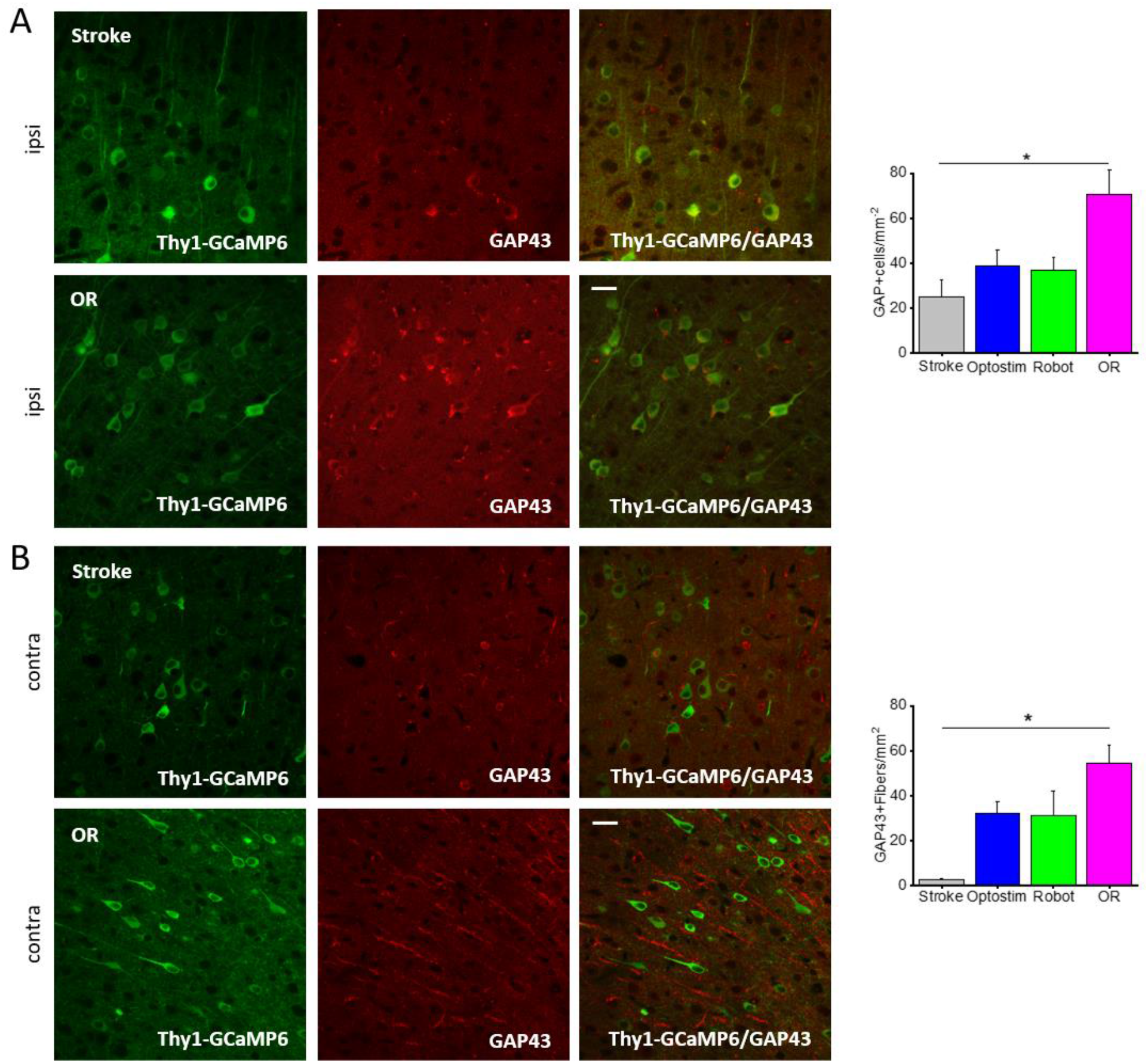
*Ex vivo* evaluation highlights the restoration of a new excitatory/inhibitory balance and the intervention of plasticizing factors in OR mice: (A) Left: Transgenic expression of GCaMP6f under Thy1 promoter (green) and representative GAP43 immunostaining (red) of a coronal section (100μm thick) of a Stroke and OR mouse. Right: quantification of GAP43+ cells in the peri-infarct area in Stroke and OR groups (Stroke= 25,0 ± 7,2; OR= 70,7 ± 10,7; *p<0.05 based on one-way ANOVA followed by Tukey’s correction: pStroke-OR= 0,011). (B) Left: transgenic expression of GCaMP6f under Thy1 promoter (green) and representative GAP43 immunostaining (red) of CL labeled fibers. Right: quantification of GAP43+ fibers in the CL hemisphere in Stroke and OR mouse (Stroke= 2,4 ± 0,4; OR= 54,4 ± 8,3; *p<0.05 based on one-way ANOVA followed by Tukey’s correction: pStroke-OR= 0,006); nStroke=4; nOptostim=4; nRobot=3; nOR=4.

## Discussion

The present study aimed at understanding the neuronal correlate of generalized functional recovery. We analyzed the motor-evoked distributed cortical activity in several rehabilitation paradigms. Among the treatments, both the optogenetic stimulation of perilesional excitatory neurons and the combined therapy with optogenetic stimulation and motor training of the paretic forelimb led to generalized recovery. However, the novel combined rehabilitation paradigm validated here led to the generalized behavioral recovery of forelimb functionality, significantly faster than optogenetic stimulation alone. Moreover the combinatory treatment induces the restoration of spatio-temporal features of cortical activity to pre-stroke levels. Based on these results our hypothesis is that generalized recovery is supported by the establishment of a new excitatory/inhibitory balance between hemispheres, revealed by the augmented cortical motor response flanked by increased expression of PV+ neurons in the peri-infarct cortex.

We first assessed the consequences of the optogenetic treatment consisting of 4 weeks of daily perilesional stimulation. Previous works demonstrated the efficacy of post-stroke optogenetic stimulation on both the peri-infarct cortex^14,15^ and striatum^43^ in promoting the recovery of forepaw sensory-motor abilities. In agreement with these findings, we reveald a remarkable improvement of forelimb functionality after 4 weeks of optogenetic stimulation of the peri-lesioned excitatory neurons. Our data also agree with the observation by Tennant and colleagues that even if optogenetic rehabilitative treatment enhanced the restoration of somatosensory cortical circuit function, the cortical area responsive to optogenetic stimulation and the peak of motor-evoked cortical activity after 4 weeks of treatment were not fully recovered to pre-stroke levels. Accordingly, our observations indicate that our optogenetic treatment *per se* does not recover pre-stroke spatio-temporal features of motor-evoked cortical activation. Indeed, a diffuse activation involving regions across the entire lesioned hemisphere is observed during active pulling of the paretic forelimb. Alongside these spread motor representations, motor-evoked cortical activation profiles have lower amplitude compared to pre-stroke conditions. Thus, our results support the hypothesis that optogenetic stimulation of excitatory spared neurons counteracts the increased excitability of the contralesional M1, thus balancing the excessive inhibitory drive onto the ipsilesional cortex. Taken together these results highlight that the restoration of pre-stroke features of cortical activity is not an essential requisite to achieve the recovery of forelimb functionality.

On the other hand, although Robot mice did not achieve a generalized recovery, their motor-evoked map is segregated to the sensorimotor regions, similarly to healthy mice (Sham group). Nevertheless, longitudinal motor training alone did not restore pre-stroke features of calcium transient. These results confirm that motor training alone promotes a task-specific motor improvement as previously shown by Spalletti and collaborators^35^, which is allegedly associated to the stabilization of motor representation^44^.

Finally, the combined rehabilitative treatment triggered a synergic effect that connects behavioral improvement to recovery of pre-stroke motor-evoked cortical activation. Indeed, the combination of cortical stimulation and motor training induces a fast restoration of the forelimb function towards healthy conditions, as measured via asymmetry index analysis in the Schallert test. At the same time, flanking the stimulation with repeated exercise leads to both confinement of motor representations and restoration of temporal features of motor-evoked calcium transient in the peri-lesioned cortex, towards the pre-stroke condition. The strong involvement of spared neurons in the secondary motor cortex revealed in the motor representation of OR mice could play a leading role in regaining generalized recovery. Our results highlight the synergic effect of the combined rehabilitation since counterbalancing the contralesional inhibition of peri-infarct cortex by optogenetic stimulation could enable the stabilization of spared circuitry achieved by longitudinal motor training.

We showed an increase in motor-evoked cortical activation mediated by excitatory cells in the peri-infarct area of OR mice compared to single treatments (Optostim and Robot groups) and spontaneous recovery (Stroke group). Together with the higher levels of parvalbumin expression, these results suggest that the combined treatment might promote the restoration of an excitatory/inhibitory balance in the peri-infarct cortex similar to pre-stroke conditions. Moreover, previous studies demonstrated that modulation in parvalbumin expression is associated to achievement of skilled ability in motor task^45^. Our results are in line with what observed by Swanson and Maffei, who shows that PV expression is stabilized to higher expressions level once motor performance saturated. In our study, the small increment of PV^+^ cells observed in Robot group is strongly enhanced in OR mice. We think that the combination of motor training with optogenetic stimulation further stimulate the inhibitory activity in the perilesional cortex.

The behavioral improvement is supported by the long-lasting expression of growth-associated factors in the perilesional area, as demonstrated by the increased number of cells expressing GAP43. It was previously demonstrated that this growth-associated factor promotes cortical remapping^3^, axonal sprouting ^3,36,37^, and dendritogenesis^41,42^ after stroke. This result extends previous findings by Cheng and collaborators showing that perilesional optogenetic stimulation after stroke promotes a global increase in GAP43 expression involving both hemispheres^14^. Also, our findings are in agreement with Spalletti and collaborators^35^ showing that a combined rehabilitative treatment leading to generalized recovery promotes an increase of plasticity markers expression. The recovery of the excitatory/inhibitory balance could be further supported by the axonal sprouting in the contralesional cortex, where a substantial number of GAP43^+^ fibers were detected, particularly in combined rehabilitated mice. The increment of GAP43^+^ cells in the peri-infarct area, in combination with GAP43^+^ axons in the contralesional region, suggests the arise of transcallosal axonal sprouting from the ipsilesional hemisphere.

One of the main results of the present study is that restoration of generalized motor function can occur without recovery of spatiotemporal features of motor-evoked cortical activation that characterize pre-stroke mice, as in Optostim mice. On the other hand, segregation of motor representation is not necessarily entangled with generalized recovery, as in Robot mice. These findings suggest that rehabilitation strategies for generalized recovery might not essentially aim at the restoration of pre-stroke features of motor-evoked activity. Nevertheless, the reestablishment of pre-stroke activation transients was a distinguished feature of the most efficient therapeutic approach, the combined therapy. In accordance with our previous findings^29^ on a different combined rehabilitation paradigm, this study supports the hypothesis that the restoration of pre-stroke temporal features could be an important biomarker of the functional recovery.

To conclude, we developed a post-stroke rehabilitation therapy that exploits the synergic effect of peri-infarct optogenetic excitation and repetitive motor training to promote the stabilization of a new excitatory/inhibitory balance between hemispheres resulting in the recovery of generalized functionality and spatio-temporal features of motor-evoked calcium transient. This study highlights how the outcome of rehabilitation therapies depends on a balance between stabilization of peri-infarct circuitry and fostering of cortical plasticity.

## Acknowledgments

We thank K. Deisseroth for opsin plasmids. We thank Matteo Caleo, Cristina Spalletti and Claudia Alia for very useful discussion about the manuscript.

## Author Contributions

E.C., A.L.A.M. and F.S.P. conceived the study. E.C., A.L.A.M. and F.C. performed experiments. E.C., A.S., G.d.V., M.P., processed data. F.S.P., T.P. and S.M. obtained funding support. E.C. and A.L.A.M. wrote the paper. All authors approved the paper.

## Data availability statement

All data that support the findings of this study are available upon reasonable request.

