## Supplementary Material for "Restoration of motor-evoked cortical activity is a distinguishing feature of the most effective rehabilitation therapy after stroke"

*Supplementary Materials*

**Mice**

All procedures involving mice were performed in accordance with the regulations of the Italian Ministry of Health authorization n. 871/2018. Mice were housed in clear plastic cages under a 12 h light/dark cycle and were given ad libitum access to water and food. We used a transgenic mouse line, C57BL/6J-Tg(Thy1GCaMP6f)GP5.17Dkim/J, from Jackson Laboratories (Bar Harbor, Maine USA). To compute the number of subjects per group we conducted a power analysis considering a one way ANOVA (fixed effects, omnibus, one-way) with an effect size $f = 1.0$, $\alpha = 0.05$ and Power = 0.8 (software used G * Power, version 3.1.9.2, Franz Faul, University of Kiel, Germany). The software calculated that the number of 4 animals per group was enough to have a power of ≈ 80. Mice were identified by earmarks and numbered accordingly. Animals were randomly divided into 5 groups and distributed as follows: Sham n=6; Stroke n=4; Optostim n=4; Robot= 7; Optostim+Robot (abbreviated in OR) n= 4. Each group contained comparable numbers of male and female mice (weighing approximately 25g). The age of mice (ranging from 6 to 8 months old) was consistent between the groups.

**Channelrhodopsin-2 (ChR2) injection**

All surgical procedures were performed under isoflurane anesthesia (3% induction, 1.5% maintenance, in 1L/min oxygen). Body temperature was maintained at 37°C and mice were monitored using respiratory rate and toe pinch throughout the procedure. The animals were placed into a stereotaxic apparatus (Stoelting, Wheat Lane, Wood Dale, IL 60191). The skin over the skull was cut and the periosteum was removed with a blade. We used a dental drill to create a small craniotomy over somatosensory cortex, which was identified by stereotaxic coordinates. We injected 0.5 μL of AAV9-CaMKIIa-hChR2(H134R)-mCherry (2.48*1013 GC/mL) 600 μm deep inside the cortex at (i) -0.75 anteroposterior, +1.75 mediolateral.

We longitudinally evaluated ChR2 expression along the weeks starting 5 days after the intracortical injection. As shown in the Supplementary Figure 1 C and D, ChR2 expression was assessed *in vivo* by acquiring through-skull fluorescence images of the reporter mCherry, which was cotransfected by our viral vector AAV9-CaMKIIa-hChR2(H134R)-mCherry. mCherry fluorescence was revealed on a wide fluorescent region of the cortex around the injection site from the first imaging session (day 5 post injection) in all transfected mice (the black spot on the top left corner of the image labels bregma). At the end of the rehabilitative period, 30 days after the injection, ChR2 expression is evenly distributed across the cortex layers and it extends for 600 μm in the rostro-caudal direction.

In Stroke, Optostim, Robot and OR mice this surgery was followed by the stroke induction. Robot mice were injected with ChR2 though the animals will not optogenetically stimulated. We applied this precaution not to underestimate a possible effect of LED excitation of ChR2 expressing neurons during calcium imaging. In Sham mice a cover glass and an aluminum head-post were attached to the skull using transparent dental cement (Super Bond, C&S), then the animals were placed in a heated cage (temperature 38°) until they fully recovered. Given the design of the experiment, no blind approach was applied during surgery. Channel rhodopsin injections were performed during the same surgical session in which the photothrombotic lesions were generated.

**Photothrombotic lesion**

The primary motor cortex (M1) was identified (stereotaxic coordinates +1,75 lateral, +0.5 rostral from bregma). Five minutes after intraperitoneal injection of Rose Bengal (0.2 ml, 10 mg/ml solution in Phosphate Buffer Saline (PBS); Sigma Aldrich, St. Louis, Missouri, USA), white light from an LED lamp (CL 6000 LED, Carl Zeiss Microscopy, Oberkochen, Germany) was focused with a 20X objective (EC Plan Neofluar NA 0.5, Carl Zeiss Microscopy, Oberkochen, Germany) and used to illuminate the M1 for 15 min to induce unilateral stroke in the right hemisphere. Sham mice were injected with 0.2 mL of saline and then illuminate as the others. A cover glass and an aluminum head-post were attached to the skull using transparent dental cement (Super Bond, C&S). We waited at least 4-5 days after the surgery for the mice to recover before the first imaging session. After the last imaging session, all animals were perfused first with 20-30 mL of 0.01 M PBS (pH 7.6) and then with 150 mL of Paraformaldehyde 4% (PFA, Aldrich, St. Louis, Missouri, USA).

**Robotic rehabilitation**

Mice were allowed to become accustomed to the apparatus before the first imaging session so that they became acquainted with the new environment. The animals were trained by means of the M-Platform, which is a robotic system that allows mice to perform a retraction movement of their left forelimb (36). Briefly, the M-Platform is composed of a linear actuator, a 6-axis load cell, a precision linear slide with an adjustable friction system and a custom-designed handle that is fastened to the left wrist of the mouse. The handle is screwed onto the load cell, which permits a complete transfer of the forces applied by the animal to the sensor during the training session. Each training session was divided into "trials" that were repeated sequentially and consisted of 5 consecutive steps. First, the linear actuator moved the handle forward and extended the mouse left forelimb by 10 mm (full upper extremity extension). Next, the actuator quickly decoupled from the slide and a tone lasting 0.5 s informed the mouse that it should initiate the task. If the animal was able to overcome the static friction (approximately 0.2 N), it voluntarily pulled the handle back by retracting its forelimb (i.e. forelimb flexion back to the starting position). Upon successful completion of the task, a second tone that lasted 1 sec was emitted and the animal was given access to a liquid reward, i.e. 10 µl of sweetened condensed milk, before starting a new cycle. To detect the movement of the wrist of the animal in the low-light condition of the experiment, an infrared (IR) emitter was placed on the linear slide, and rigidly connected to the load cell and thus to the animal's wrist. Slide displacement was recorded by an IR camera (EXIS WEBCAM #17003, Trust) that was placed perpendicular to the antero-posterior axis of the movement. Position and speed signals were subsequently extracted from the video recordings and synchronized with the force signals recorded by the load cell (sampling frequency = 100 Hz). To adjust the friction of the device, a calibration procedure was performed by connecting the actuator with the slide using a rigid component. This component mimics the effect of the retraction movements performed by the animal, reproducing the applied forces in direction and point of application. The training consisted of 15 cycles of passive extension of the affected forelimb followed by its active retraction triggered by the acoustic cue. All groups performed at least one week (5 sessions) of daily training, starting 26 days after injury for Stroke and Optostim mice, 5 days after stroke for Robot and OR groups and after the surgery for Sham animals. During the last week of optogenetic stimulation of Optostim and Stroke mice and the entire rehabilitative period for OR mice the robotic training is followed by the optogenetic stimulation.

**Optogenetic stimulation**

Awake head-fixed mice were placed under the wide-field microscope to perform daily session of optogenetic stimulation. A blue 473 nm laser is used to deliver 5 Hz, 10ms light pulses similar to Tennant 2017 15. The laser power used, ranging from 0.2 to 0.8 mW, was lower than the one necessary to elicit movements of the affected forelimb during cortical optogenetic stimulation. Furthermore, the laser power is adjusted during the rehabilitation period, according to the increment of the transfected area and the progressive lowering of stimulation threshold over the weeks. The system is provided with a random-access scanning head, developed using two orthogonally mounted acousto-optical deflectors (DTSXY400, AA Opto-Electronic). The acousto-optic deflectors rapidly scan lines with a commutation time ~5 μs between a line and the next. After scanning the desired shape (in this case a cross) in the area of the cortex of ChR2 maximum expression (approximately near the injection site -0.75 anteroposterior, +1.75

mediolateral), the acousto-optic deflectors returned to the initial position and repeated the cycle for the total illumination time. The stimulation protocol consists of 3 successive 30 s stimulation daily separated by 1 min rest intervals. All animals (Stroke, Optostim and OR), except for Sham and Robot groups, were stimulated every day for 4 weeks, 5 days after photothrombosis. Optostim and OR groups are exposed to optogenetic stimulation approximately 5 minutes after the execution of the last pulling during motor rehabilitation. We stimulated Stroke mice not expressing ChR2 to evaluate possible artefacts due to repeated laser stimulation. All the experiments revealed that repeated stimulation of mouse cortex not expressing ChR2 did not affect post-stroke recovery.

**Schallert Cylinder Test**

Mice were placed in a Plexiglas cylinder (7,5 cm diameter, 16 cm height) and recorded for five minutes by a webcam placed below the cylinder, after 2 minutes of acclimatization. Videos were analyzed frame by frame and the spontaneous use of both forelimbs was assessed during exploration of the walls, by counting the number of contacts performed by the paws of the animal. For each wall exploration, the last paw that left and the first paw that contacted the wall or the ground were assessed. The analysis was conducted in blind. In order to quantify forelimb-use asymmetry displayed by the animal, an Asymmetry index was computed, according to Lai et al. 2015[46].

**Wide-field fluorescence microscope**

The custom-made wide-field imaging setup was equipped with two excitation sources allowing imaging of GCaMP6f fluorescence and light-stimulation of ChR2. For imaging of GCaMP6f fluorescence, a 505 nm LED (M505L3 Thorlabs, New Jersey, United States) light was deflected by a dichroic filter (DC FF 495-DI02 Semrock, Rochester, New York USA) on the objective (2.5x EC Plan Neofluar, NA 0.085, Carl Zeiss Microscopy, Oberkochen, Germany). A 3D motorized platform (M-229 for xy plane, M-126 for z-axis movement; Physik Instrumente,

Karlsruhe, Germany) allowed sample displacement. The fluorescence signal was selected by a band-pass filter (525/50 Semrock, Rochester, New York USA). Then a 20X objective (LD Plan Neofluar, 20x/0.4 M27, Carl Zeiss Microscopy, Oberkochen, Germany) was used to demagnify the image onto a 100X100 $pxl^2$ area of the sCMOS camera sensor (OrcaFLASH 4.0, Hamamatsu Photonics, NJ, USA). Images (5.2 x 5.2 $mm^2$, pixel size 52 µm) were acquired at 25 Hz. To perform optogenetic stimulation of ChR2, a 473 nm continuous wavelength (CW) laser (OBIS 473nm LX 75mW, Coherent, Santa Clara, California, United States) was overlaid on the imaging path using a second dichroic beam splitter (FF484-Fdi01-25x36, Semrock, Rochester, New York USA) and scanned with two orthogonally-mounted acousto-optical deflectors as previously described (DTSXY400, AA Opto-Electronic, Orsay France).

**Image analysis**

Analysis of the fluorescence image stacks were previously reported in Allegra et al. 2019. We aimed to characterize activation maps and calcium transient critical features, focusing those transients occurring in the time-window of retraction movement. For each stack (FluoSt) a median time series of GCaMP6f fluorescence signal (mF) was extracted, where the value of mF at each ith time point corresponded to the median value computed on all the pixels of the i-th frame of FluoSt. The mF was then oversampled and synchronized to the 100 Hz force and position signals. The mF was used to define a GCaMP6f fluorescence signal baseline F0, which was identified by the concomitant absence of fluorescence and force signal deflections. F0 was selected within a 2.5 ± 0.7 s interval (I) of 62 frames where the fluorescence signal was below 1 standard deviation (STD) of the whole mF signal and the corresponding force signal showed a value below 1 STD of the whole recorded force signal. The onset of each force peak was used as a reference time point to select a sequence of 60 frames (2.4 s, where 0.4 s preceded the force peak) from FluoSt. All sequences were visually checked to exclude possible spurious activation (e.g., early activation or no activation) from the analysis. All the selected sequences (Seqs) of the animal An on day d were compiled, defining a stack of Seqs, to compute the Summed Intensity Projection for the An at d (SIPAn d). The most active area of the SIPAn d was then detected by thresholding the SIPAn d with a median (SIPAn d) + STD (SIPAn d) threshold value. The threshold SIPAn d (th-SIPAn d) computed for each week of training on the M-Platform (d = 1,.,4 of week W) was superimposed and the common areas, activated at least for 3 daily sessions out of 5 (60%), were labeled as ‘‘regions of interest’’ (ROIs) of the SIPAn. We also used the ROIs defined for each individual animal to identify the average ROIs among mice from the same experimental group (Figure 2D, 3D). The ROIs defined for each individual animal were further used to extract the GCaMP6f fluorescence signal corresponding to the activity of those areas. Indeed, from each frame of the FluoS, only pixels belonging to the selected ROI were considered when calculating the representative median value. Thus, a median time series, FROI, was extracted from the whole FluoS and was representative of the ROI. The fluorescence signal was normalized ((ΔFROi /F0)100%) and low-pass filtered to clean the signal from the detected heart-beat artifacts (Chebyshev filter with cutting frequency = 9 Hz). The previously detected force peaks were then used to select the GCaMP6f fluorescence peaks from the(ΔFROI/F0) signal. A time window that lasted 4 s, i.e., wnd, and was centered at the onset of the force peak, was used to delimit a part of the (ΔFROI/F0) signal, i.e., Fwnd, to identify the corresponding fluorescence peak. A fluorescence peak was defined as the part of Fwnd that overcame the value of median+3STD calculated for the whole signal(ΔFROI/F0).

**Force analysis**

To quantify the behavior of each animal on the platform we analyzed the force exerted by the mouse during the retraction phase. For each peak associated with the slide movement we extracted 4 parameters: 1) peak amplitude as its max value; 2) peak width (full width at half maximum) as the width of the peak at half the peak amplitude; 3) peak area as area of the force signal during its peak width and 4) the time to target as the time between the start of the trial and the peak associated with the last movement of the slide in the trial.

**Immunohistochemical analysis**

For immunohistochemical analysis of plasticity markers, animals were transcardially perfused with 4% paraformaldehyde. Brains were cut using a vibrating-blade vibratome (Leica, Germany) to obtain 100 µm thick coronal sections that were used for immunostaining of NeuN (1:200, Millipore, Germany), Parvalbumin (1:500, Abcam, United Kingdom) and GAP 43 (1:600, Abcam, United Kingdom). The number of Parvalbumin- and GAP43- positive neurons was analyzed using a confocal fluorescence microscope (Nikon Eclipse TE 300, Tokyo, Japan) with a Nikon Plan EPO 60× objective (NA 1.4, oil immersion Nikon, Tokyo, Japan), acquiring 212 µm wide images in the peri-infarct cortex (up to 600 µm from the damage in the caudal direction). For GAP43 and PV evaluations three sections per animal were analyzed. In each section the 3 images (212x212 $\mu m^{2}$) including layer II/III and V of the cortex (from the longitudinal fissure up to 2 mm in the mediolateral direction) were acquired. We excluded the corpus callosum from the quantification of the fibers. The single fibers revealed in the contralesional hemisphere were measured in the same brain section used for the ipsilesional oneTo evaluate the extension of the lesion we use a 10× objective (C-Apochromat, 0.45 NA, Carl Zeiss Microscopy, Oberkochen, Germany field of view 1.3 mm). Then, the stroke volume for each animal was calculated by summing up all damaged areas and multiplying the number by section thickness and by 3 (the spacing factor). A total volume in $mm^3$ is given as the mean ± standard error of all analyzed animals (n=15). The experimenter was blind to the experimental group of the samples.

**Reagent identification**

| REAGENT or RESOURCE | SOURCE | IDENTIFIER |
|---|---|---|
| Antibodies | | |

| | | |
|---|---|---|
| NeuN | Merck | ABN78 |
| GAP43 | Abcam | Ab16053 |
| PV | Abcam | Ab11427 |
| Alexa Fluor 568 Goat anti Rabbit | Abcam | Ab175471 |
| Virus Strains | | |
| AAV9-CaMKII-ChR2-mCherry | DBA Italia | VB4411 |
| Chemicals, Peptides, and Recombinant Proteins | | |
| Zoletil | Virbac | 101580025 |
| Xylazine | Dechra | 103595017 |
| Rose Bengal | Sigma | 330000 |
| Phosphate Buffer Saline | Sigma | P-4417 |
| Lidocaine 2% | Zoetis Italia srl | 100319019 |
| Dexamethasone | MSD | 101866034 |
| Paraformaldehyde | Sigma | 158127 |
| Experimental Models: Organisms/Strains | | |
| Mouse: C57BL/6J-Tg(Thy1GCaMP6f)GP5 | The Jackson Laboratories | 025393 |
| Software and Algorithms | | |
| ImageJ | Schneider et al., 2012 | https://imagej.nih.gov/ij/ |

**Supplemental Figures**

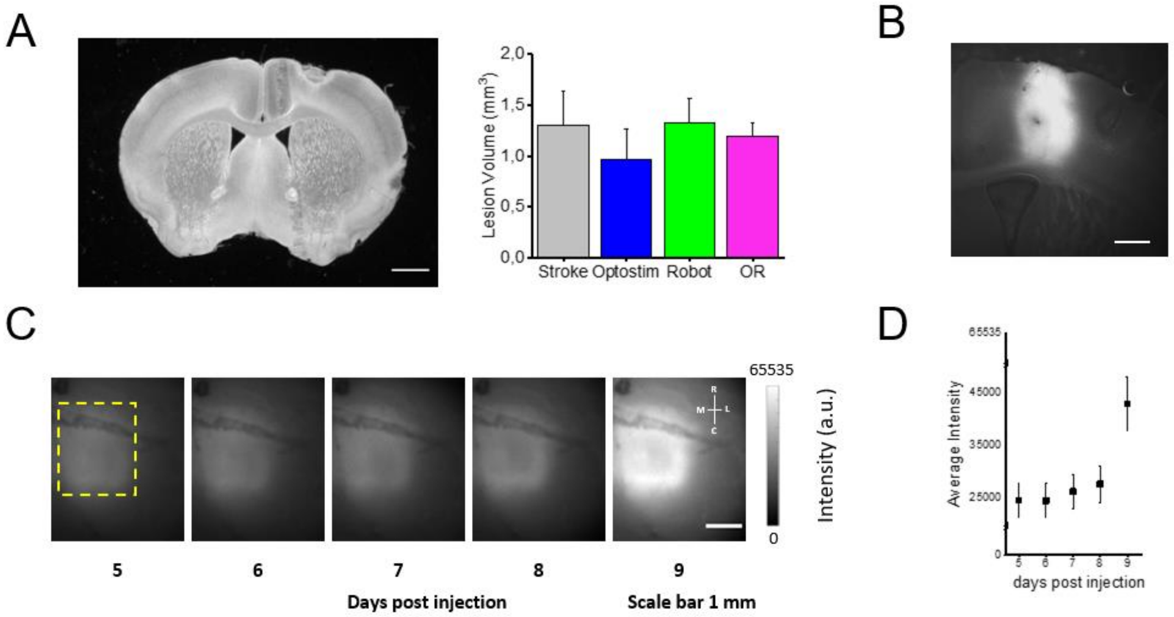


**Supplementary Figure 1.** (A) Coronal section of a mouse brain 30 days after photothrombosis, 1 mm scale bar. The right panel shows the quantification of the lesioned area in the experimental groups (average ± SEM): Stroke=1,31 ± 0,32 mm$^2$; Optostim=0,96 ± 0,3 mm$^2$; Robot=1,33 ± 0,24 mm$^2$; OR=1,2 ± 0,13 mm$^2$. (B) Coronal section of a mouse brain transfected with ChR2 30 days after the injection, 0.5 mm scale bar. (C) Longitudinal imaging of mCherry fluorescence to evaluate ChR2 expression in the peri-infarct area during the first week of stimulation (starting 5 days after the injection) (D) Quantification of average intensity (Average ± SEM) of mCherry fluorescence during the first week of stimulation (d5 = 24645.0 ± 3163.8; d6 = 24594.2 ± 3216.3; 26302.1 ± 3240.1; d8 = 27674.9 ± 3447.2; d9 =42896.1 ± 5171.7 ;).

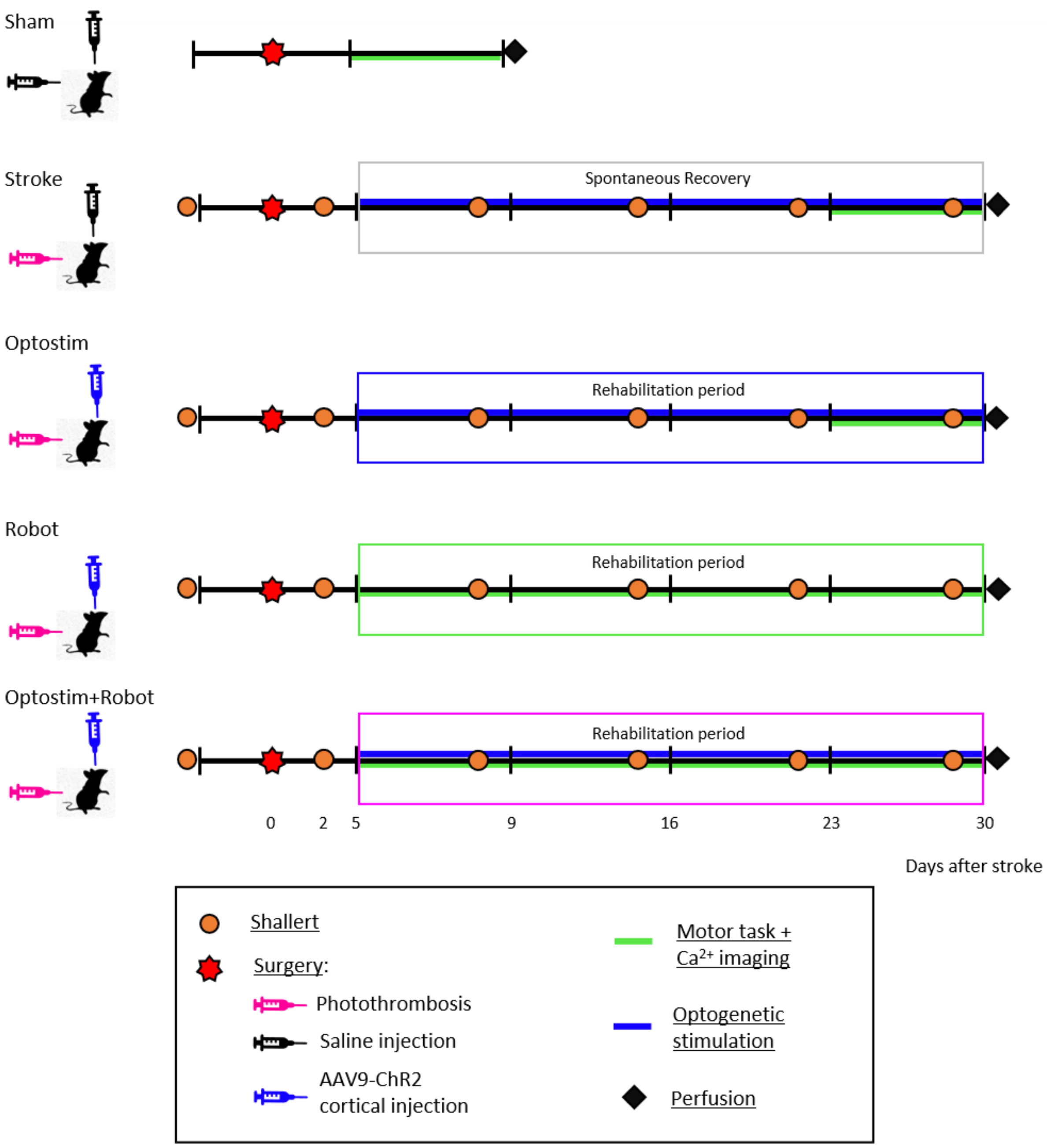


**Supplementary Figure 2.** Schematic of the treatment protocols. Baseline performances in behavioral tests (orange dot) were assessed for all groups before the surgery and then once a week up to 30 days after the lesion. The last day of treatment mice were perfused (black rhombus).

Sham: ip injection of saline (no stroke), intra cortical injection of saline (no ChR2 expression). 5 days of motor assessment on the M-Platform.

Stroke: stroke, ip injection of Rosebengal, intra cortical injection of saline (no ChR2 expression). 4 weeks of peri-infarct laser stimulation. 5 days of motor assessment on the M-Platform, 26 days after the lesion.

Optostim: stroke, ip injection of Rosebengal, intra cortical injection of AAV9 (ChR2 expression). 4 weeks of peri-infarct laser stimulation. 5 days of motor assessment on the M-Platform, 26 days after the lesion.

Robot: stroke, ip injection of Rosebengal, intra cortical injection of AAV9 (ChR2 expression, no laser stimulation). 4 weeks of motor rehabilitation on the M-Platform, starting 5 days after the lesion.

Optostim+Robot (OR): stroke, ip injection of Rosebengal, intra cortical injection of AAV9 (ChR2 expression). 4 weeks of motor rehabilitation on the M-Platform and laser stimulation, starting 5 days after the lesion.

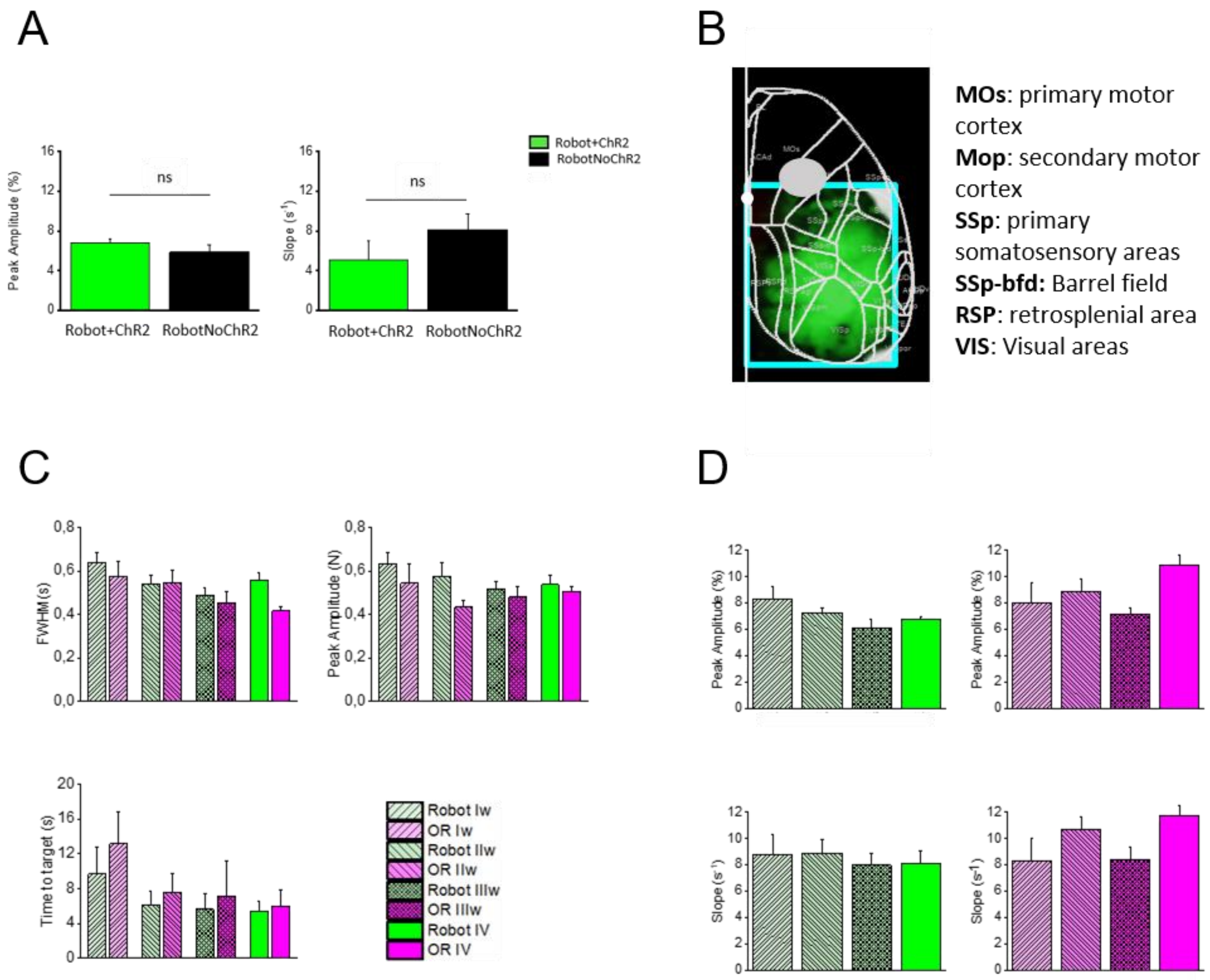


**Supplementary Figure 3.** (A) The graphs show calcium transient analysis for Robot mice expressing ChR2 (n=5) in the peri-infarct cortex or not (n=3). The left panel shows the Peak Amplitude (average ± SEM) Robot+ChR2 = 6.7 ± 0.3%; RobotNoChR2 = 5.8 ± 0.8%. The right panel shows the Slope (average ± SEM) Robot+ChR2= 8.1 ± 0.9 $s^{-1}$; RobotNoChR2= 5.1 ± 0.8 $s^{-1}$. (B) Graphical representation of functional regions of the cortex acquired in the imaging area (cyan square) including primary and secondary motor area, primary somatosensory and barrel field, retrosplenial and primary visual areas. The gray circle on the M1 region highlights the location and approximate extent of the lesion. White dot indicates bregma. (C) Force analysis during pulling task (average ± SEM). Left upper panel shows the Full Width Half Maximum of the force peak (Robot_Iw= 0,64 ± 0,05; OR_Iw= 0,58 ± 0,07; Robot_IIw= 0,54 ± 0,04; OR_IIw= 0,55 ± 0,06; Robot_IIIw= 0,49 ± 0,03; OR_IIIw= 0,45 ±

0,05; Robot_IVw= 0,56 ± 0,04; OR_IVw= 0,42 ± 0,02). Right upper panel show the Peak Amplitude (Robot_Iw= 0,63 ± 0,05; OR_Iw= 0,54 ± 0,18; Robot_IIw= 0,58 ± 0,06; OR_IIw= 0,43 ± 0,06; Robot_IIIw= 0,52 ± 0,03; OR_IIIw= 0,48 ± 0,04; Robot_IVw= 0,54 ± 0,04; OR_IVw= 0,5 ± 0,04) and the lower panel shows the Time to Target (Robot_Iw= 9,7 ± 3,2; OR_Iw= 13,2 ± 3,6; Robot_IIw= 6,1 ± 1,6; OR_IIw= 7,6 ± 2,1; Robot_IIIw= 5,6 ± 1,8; OR_IIIw= 7,1 ± 4,1; Robot_IVw= 5,4 ± 1,2; OR_IVw= 6,0± 1,8). (D) Longitudinal analysis of cortical activation profiles over the 4 weeks for Robot (Amplitude: Iw= 8.3 ± 1.0; IIw= 7.3± 0.3; III= 6.1± 0.6; IV= 6.7 ±0.3; Slope: Iw= 8.8± 1.5; IIw= 8.9 ± 1.0; III= 8.0 ± 0.9; IV= 8.1 ± 0.9) and OR groups (Amplitude: Iw=8.1 ± 1,4; IIw= 8.9 ± 0.9; III= 7.1 ± 0.5; IV= 10.9 ± 0.7; Slope: Iw= 8.3 ± 1.6; IIw= 10.7 ± 0.9; III= 8.4 ± 0.9; IV= 11,8 ± 0.7) same groups legend as in panel C.

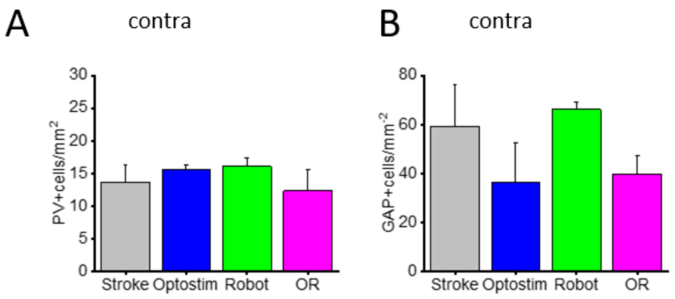


**Supplementary Figure 4.** (A) Quantification of PV+ cells in the contralesional hemisphere (Stroke= 13,6 ± 2,6; Optostim= 15,5 ±0, 7; Robot= 16,0 ± 1,4; OR= 12,2 ± 3,3). (B) Quantification of GAP43+ cells in the contralesional hemisphere (Stroke= 59,1 ± 16,9; Optostim= 36,5 ± 16,1; Robot= 66,1 ± 3,2; OR=39,6 ± 7,6).
